# Tool and hand adaptation and localization in immersive virtual reality

**DOI:** 10.1101/2025.09.02.673854

**Authors:** M.S. Khan, S. Modchalingam, A. King, B. M. ’t Hart, D. Y. P. Henriques

## Abstract

The human brain readily adapts movements to achieve motor goals. Most visuomotor adaptation studies use planar reaching with a cursor, where misaligned feedback leads to compensatory adjustments, reach aftereffects, and shifts in perceived hand location. Whether these effects extend to more natural settings and tool use remains unclear. In the Hand Experiment, we showed that in immersive virtual reality (VR), adaptation to 30° and 60° visuomotor rotations produced robust reach aftereffects and the expected shifts in hand localization. In the Pen Experiment, we extended this paradigm to a familiar hand-held tool, assessing localization of both the tool tip and the hand. Adaptation with the pen induced comparable or greater recalibration effects than with the hand-cursor, including shifts in both perceived tool and hand position. These findings demonstrate that visuomotor adaptation in immersive VR generalizes beyond cursor-based tasks, revealing how the sensorimotor system recalibrates internal representations of both the body and tools in realistic 3D environments.

**Author summary for PLOS ONE:** This study shows that when people adapt their movements in virtual reality, the brain recalibrates not only the sensed position of the hand but also of familiar tools, highlighting how we update body and tool representations in everyday-like environments.

## Introduction

Humans continuously adjust their movements to maintain accurate performance despite changes in the body, environment, or technology. This sensorimotor adaptation supports everyday functions such as compensating for fatigue or injury, learning to use new tools, and interacting with novel interfaces, including immersive virtual reality. As these technologies become more prevalent, understanding how the brain adapts to altered visuomotor relationships is increasingly important.

Sensorimotor adaptation is a form of short-term motor learning that unfolds over minutes to hours and engages both implicit and explicit processes. It is often studied using visuomotor rotation paradigms, where visual feedback of the hand—typically shown as a cursor—is rotated relative to actual movement. This mismatch between intended and perceived outcomes prompts the brain to adjust motor output. For example, with a 30° counterclockwise cursor rotation, participants must move their hand 30° clockwise to reach the target, and typically adapt within about 20 trials. When the perturbation is removed, participants continue reaching in the adapted direction, producing aftereffects that reflect implicit recalibration of motor commands ([1–8]). These aftereffects occur even when participants are told the perturbation is gone and instructed not to compensate, and they tend to saturate at ∼15° regardless of rotation size [5,9,10].

In addition to altering motor output, visuomotor adaptation also shifts internal estimates of hand position, which are critical for accurate motor planning and control. These estimates integrate visual inputs, proprioceptive signals from muscles, joints, and skin, and efference copies of motor commands that are used to predict sensory consequences of planned movements. Together these signals support smooth, coordinated movements and quick corrections. Following adaptation to altered visual feedback, people often experience proprioceptive recalibration—a shift in the perceived movement direction of their adapted end effector[9,11–17]. In our example adaptation to a 30° visuomotor rotation, these shifts are usually smaller than reach aftereffects (typically ∼ 6°) but are robust and emerge quickly, within several trials [11,16–20].

Proprioceptive recalibration occurs even when the misalignment is consciously recognized, reinforcing its implicit nature [9,10,21–23]. Localization shifts can be measured using active localization (self-generated placement of the trained hand) or passive localization (robot-controlled placement of the trained hand). Active localization tends to yield slightly larger shifts, likely reflecting a combination of proprioceptive and efferent signals, whereas passive localization isolates proprioceptive signals [9,15,22,24,25].

Most of this work has been conducted in 2D screen-based setups using cursors. Yet in the real world, we interact with objects in 3D environments, frequently often via tools rather than directly with our hands. It remains unclear whether the same adaptation and proprioceptive recalibration processes extend to more naturalistic contexts. Immersive virtual reality (VR) offers a unique platform to test this question because it can simulate realistic environments while maintaining precise experimental control. Immersive VR is increasingly used in motor learning and rehabilitation due to its safety, cost-effectiveness, and ecological validity [26,27]. Although visuomotor adaptation and aftereffects have been demonstrated in VR [28–34], some studies suggest that increased cognitive load in VR could reduce adaptation compared to traditional setups [29], while others report similar adaptation [30]. This study addresses this gap by examining whether adaptation and proprioceptive recalibration observed in traditional 2D paradigms are preserved when interacting with tools in immersive VR, thereby providing insight into the generalizability of these processes to more naturalistic settings.

Another key aspect of natural motor behavior is tool use, where tools such as pens, bats, or utensils become integrated into the body schema through a process known as tool embodiment [35–37]. Neurophysiological evidence shows that tool use can expand receptive fields in the intraparietal cortex, effectively extending the brain’s representation of peripersonal space to include the tool [35,37–43]. In visuomotor adaptation tasks, changes in reach direction or amplitude resemble the adjustments required for controlling tools, particularly when the tool alters the spatial relationship between the hand and its effect on the environment (e.g., Alzayat et al., 2019). However, few studies have examined whether sensory recalibration extends to tools themselves, especially in immersive VR.

In the present study, we addressed these gaps by examining visuomotor adaptation and proprioceptive recalibration in immersive VR. In the Hand Experiment, we replicated prior work from our lab showing shifts in hand localization after adaptation to 30° and 60° visuomotor rotations [9,11,16,21,25]. In the Tool Experiment, we extended this paradigm to a familiar hand- held tool—a virtual pen—whose tip movements were rotated by 30°. We then assessed whether adaptation recalibrated both the perceived tool-tip location and the hand itself (in separate trials).

Together, these experiments aimed to determine whether visuomotor adaptation in immersive VR produces comparable motor aftereffects and proprioceptive recalibration to traditional 2D paradigms and whether these recalibration processes extend beyond the hand to a commonly used tool. We hypothesized that participants wielding a pen would adapt similarly in VR and show robust shifts in both reach aftereffects and localization. Additionally, we explored whether tool-tip adaptation would induce recalibration not only for the tool but also for the underlying hand. By comparing adaptation and recalibration across direct hand use and tool use in VR, we sought to clarify how the sensorimotor system flexibly updates internal representations of both the body and hand-held objects in realistic, immersive environments.

## Methods

### Participants

A total of 64 undergraduate students (mean age = 21.8 ± 5.9 years, 45 females) recruited from York University participated in the study between January 2023 and April 2024. Participants were run in at least one of the three groups: 30° hand, 60° hand and 30° pen groups. All participants reported having normal or corrected to normal vision, and gave prior, written, informed consent. Nearly all participants were right-handed, except for two that were left-handed and one ambidextrous. The procedures used in this study were approved by York University’s Human Participants Review Subcommittee.

### Experimental set-up

#### Apparatus

Participants sat on a height-adjustable chair facing a table that was at waist level. They put on a head-mounted display system (HMD: Oculus Rift Consumer Version 1; resolution 1080 by 1200 for each eye; total refresh rate 90 Hz) and held an Oculus Touch controller in each hand. All visual feedback was provided through the HMD in an immersive VR experiment space developed in Unity 3D, with trial and block schedules managed with the Unity Experiment framework [44]. The virtual scene was designed to match the physical dimensions of the table (3.5 m wide × 2 m deep) to provide consistent spatial alignment between real and virtual space (as with Modchalingam et al., 2024, 2025). A docking station was located at the edge of the table just in front of the body, along the midline (location indicated in Fig 1). A small magnet on the controller and docking station ensured that participants consistently returned the controller to the same start location at the beginning of each trial. As consistent with previous work in our lab [31,32], three Oculus Constellation sensors tracked the positions of the headset and controllers, with positional data obtained via the Meta software. For setups similar to the one used in this study, positional measurement errors are expected to be within millimeters, with errors in the non-vertical dimensions expected to be less than 1% for the movement size used in this study [45,46].

**Figure 1:**
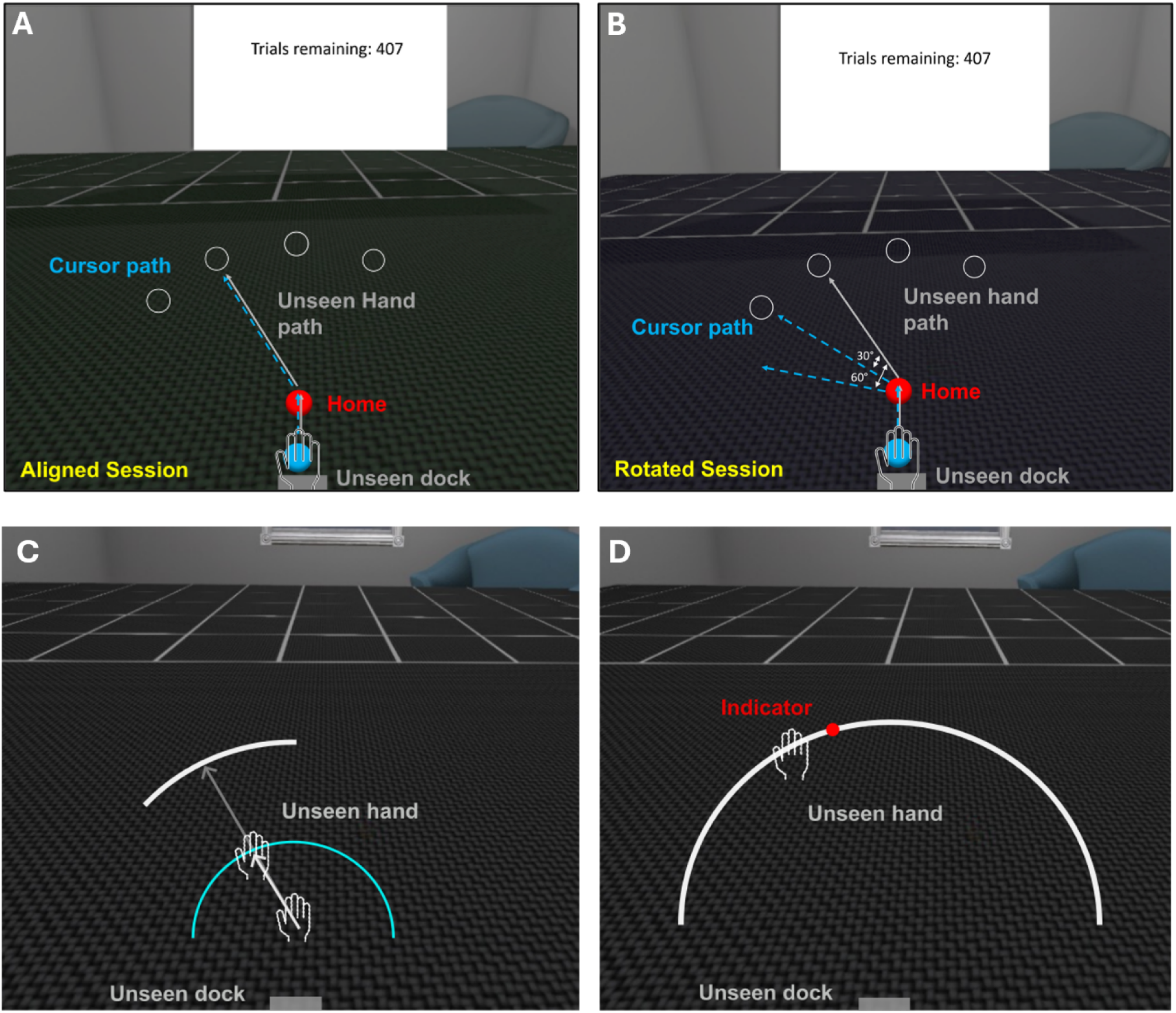
Hand experimental stimuli. **A)** In ‘Reach to Target’ tasks in the ‘Aligned’ session, the blue cursor representing the hand followed the unseen hand path (**B)** In the ‘Rotated’ session of the experiment, a 30° or 60° CCW perturbation applied to the blue cursor so that participants had to reach 30° or 60° to the right to match the cursor path to the unseen hand path. **C)** In the Hand Localization trials, participants reached out to anywhere along a white arc using their unseen right hand **(D)** Then, participants used their left controller’s joystick to move the red indicator to mark where their unseen right hand crossed the white arc. The blue arc expanded as the hand moved outward for participants to know how far they need to reach out to acquire the arc.

#### General Procedure

Participants were instructed to face forward throughout the study, with a slight downward tilt of the head to view the stimuli within the VR environment. At the start of the session, the experimenter calibrated the display by centering the visual stimuli around the location of the right-hand controller of the participant at the physical docking station. From there, participants moved out their hand-held controller to visually indicate home position, and then either to a target, or an arc (Figure 1) depending on the task, as described below. A trial counter displayed the number of remaining trials. Participants were instructed to smoothly glide their arm along the tabletop, making straight, accurate, and consistent movements to either the target or arc.

The study consisted of two types of conditions when reaching for the target: *Hand* and *Pen*. In the Hand conditions, participants reached to targets using their right index finger, represented by a blue cursor (Figure 1). In the Pen conditions, they reached using the tip of a virtual pen, with a physical pen secured to the right-hand controller to provide tactile feedback and promote realistic tool use. The virtual environment and task structure were similar across conditions. The Pen environment depicted an office scene with a desk surface, while the Hand environment used a neutral room with a dark green table. The Hand condition was conducted with both 30° and 60° cursor rotations (explained below), while the Pen condition included only the 30° rotation to allow direct comparison with the corresponding Hand group.

### Hand Experiment

#### Reach to Target tasks

At the start of each block of trials, ‘Reach to Target’ was displayed. Participants held the cursor (blue sphere, 1 cm diameter) at the central home position for 300 ms, then reached to one of four targets (red spheres, 1 cm diameter) located 12 cm away at 70°, 90°, 110°, or 130° (Figure 1). To acquire the target, participants held the cursor within 0.5 cm of its center for 300 ms. The cursor and target then disappeared, and participants returned to the dock.

In the Rotated session (Figure 1B), a 30° or 60° counterclockwise (CCW) rotation was applied to the cursor motion direction. To compensate, participants had to move 30° or 60° clockwise (CW) to acquire the target.

### No Cursor

These trials measured aftereffects following visuomotor adaptation training. At the start of this block of trials, ‘No Cursor’ was displayed. Participants reached to the same four targets without visual feedback of the cursor. A blue arc, spanning 180° and centered on the home position, tracked outward movement (blue curve in Figure 1D). Targets appeared against a dark gray surface to signal this trial type.

Participants moved their hand at least 3 cm from home and held their position for 500 ms to complete the trial. The hold was verified by ensuring that frame-to-frame hand displacement remained below 5 mm during this period. Afterward, the arc and target disappeared, and participants returned to the dock. To isolate implicit components of adaptation, participants were instructed to reach naturally and not use previously learned strategies developed that they used while reaching with the cursor.

### Localization

In the localization trials, participants estimated the location of their unseen hand. While holding the controller in their right hand, they made volitional hand movements from the home position to self-selected points along a white arc (0.5 cm thick), located 12 cm away (Figure 1C). To sample a wide range of hand or pen locations, this arc spanned 60°, with its center randomly set to 70°, 90°, 110° or 130° in polar coordinates across trials. As in the No Cursor trials, a blue arc, spanning 180° indicated how far the hand had moved. Once the hand moved 12 cm from the home position and remained still for 500 ms while the blue arc overlapped with the white arc, a 180° white arc was displayed. Using the left controller’s joystick, participants then moved a red indicator along this arc to the point where they felt their index finger had intersected it (Figure 1D). They confirmed their estimate by briefly pressing the trigger on the right controller. Once the arc and indicator disappeared, participants returned their right hand to the dock to await the next trial. This procedure was used to assess perceived hand position both before and after adaptation to the rotated cursor.

#### Schedule

In the Hand Experiment, we aimed to replicate previous findings from studies using a robot manipulandum, but within an immersive VR setup. Before the main experiment, participants completed a brief practice session (16 Hand-to-Target, 8 Localization, and 8 No Cursor trials) to familiarize themselves with the task. As shown in Figure 2A, participants completed two sessions: an Aligned session and a Rotated session. During the Aligned session, the visual cursor accurately represented the hand’s movement direction. The Aligned session was repeated twice and began with 32 Hand-to-Target trials, followed by alternating blocks of Hand Localization and additional Hand-to-Target trials. A Hand No Cursor block occurred after the final Hand-to-Target block, at the end of each repetition. Across these blocks, each target position was presented twice in a pseudo-randomized order. Participants were given short breaks between repetitions and a longer break after the Aligned session was complete, before moving on to the Rotated session. The VR headset remained on throughout the entire procedure, including breaks.

**Figure 2.**
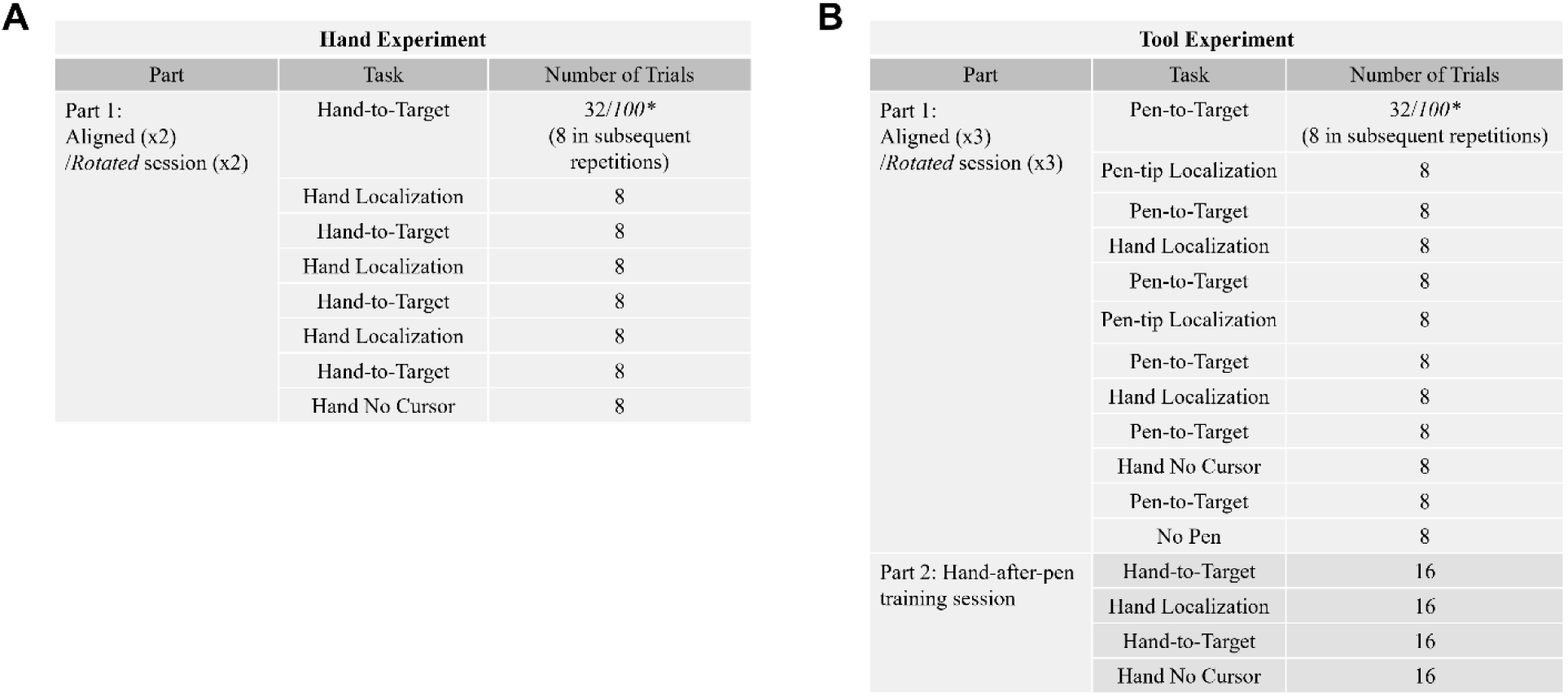
Experiment schedule for Hand and Tool Experiments. A) In the Hand Experiment, participants performed two repetitions of the aligned session and then took a mandatory short break before completing two repetitions of the rotated session. The Hand-to-Target tasks in the rotated session consisted of a 30° or 60° CCW visuomotor rotation. B) In the first part of the Tool Experiment, participants performed three repetitions of the aligned session and then, after a mandatory break, three repetitions of the rotated session. In the second part of this experiment, participants trained their hand after training with the pen. *The aligned session began with 32 trials and the rotated session with 100 trials, followed by repetitions of 8 trials each. All trials were organized into sets of four, with one reach to each target per set in random order.

Following the mandatory one-minute break, participants completed the Rotated session, in which the cursor was rotated either 30° or 60° counterclockwise for all Hand-to-Target trials (Figure 2A). The structure of the Rotated session mirrored that of the Aligned session, except the first Reach to Target block was extended from 32 to 100 trials to allow sufficient adaptation to the visuomotor perturbation. Participants were informed that the cursor would behave differently but were instructed to continue aiming directly for the target with fast, straight, and accurate reaches. This was followed by the same sequence of blocks—Localization, No Cursor, and additional Hand-to-Target trials—to assess post-adaptation changes. During No Cursor trials, participants were instructed to exclude any conscious strategy used during training so that any reach aftereffects would reflect implicit adaptation. At the end of the Rotated session, the study was complete, and participants were instructed to remove the headset, marking the end of the ∼50-minute experiment.

### Pen Experiment

The goal of the Pen experiment was to examine whether similar adaptation and localization shifts would emerge when participants reached using a familiar tool — a virtual pen (displayed on top of a physical pen). The procedure was identical to the previously described Hand Experiment, except that participants used the tip of the pen, rather than a hand-aligned cursor, to acquire targets (Figure 3).

**Figure 3:**
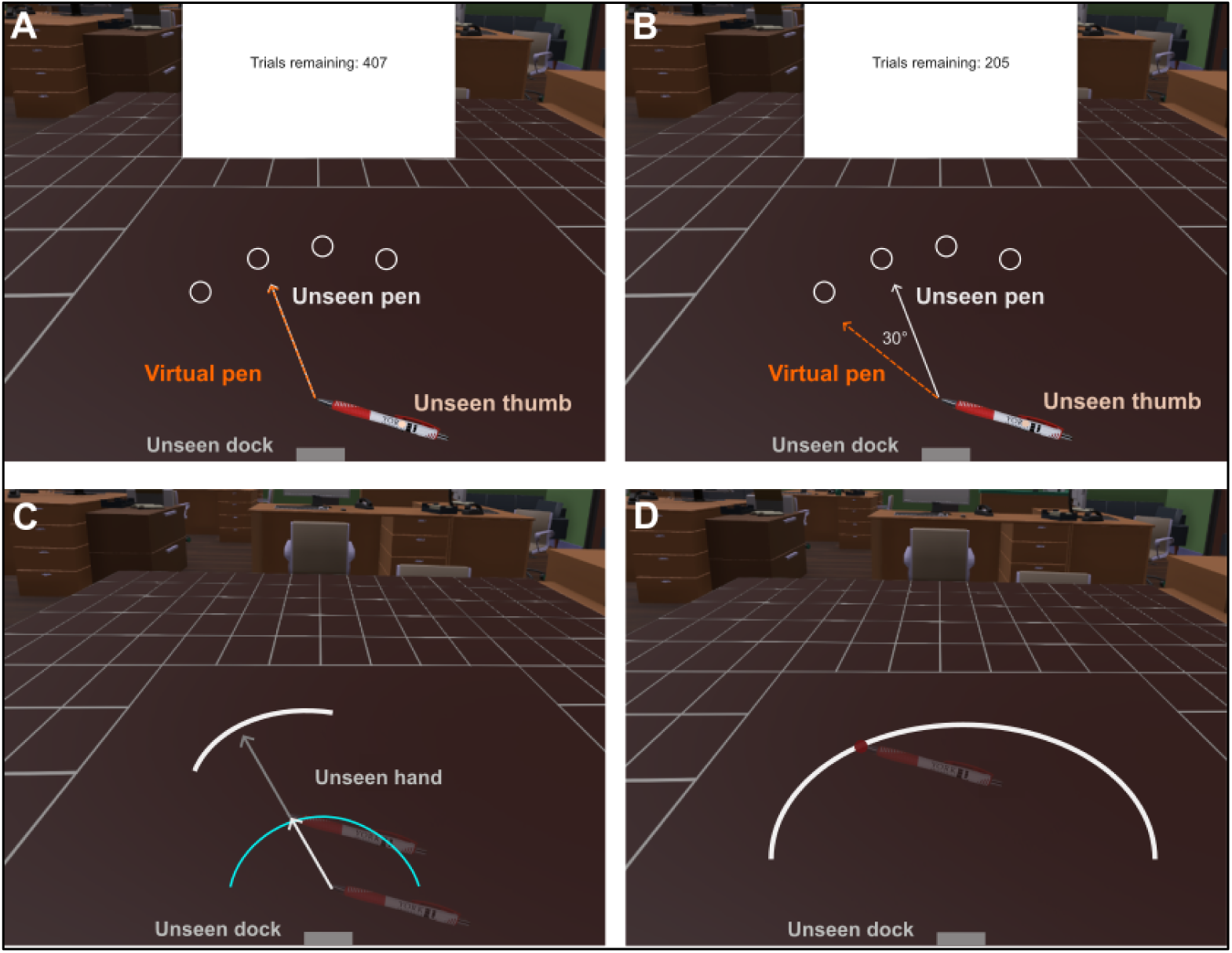
Pen experimental stimuli. **A)** In pen ‘Reach to Target’ tasks in the ‘Aligned’ session, the pen tip followed the unseen physical pen path. (**B)** In the ‘Rotated’ session of the experiment, a 30° CCW perturbation was applied to the tip of the pen so that participants had to reach 30° to the right with the physical pen to match the virtual pen path to the physical pen path. (**C)** In the pen localization task, participants reached out to anywhere along a white arc using the unseen tip of the pen. **(D)** Then, participants used their left controller’s joystick to move the red indicator to mark where the unseen tip of the pen crossed the white arc. The blue arc expanded as the pen tip moved outward for participants to know how far they need to reach out to acquire the arc.

Participants began with a practice session (16 Pen-to-Target, 8 Pen Localization, and 8 No Pen trials) to familiarize themselves with pen-based reaching. The main experiment followed the same structure as the Hand Experiment (Figure 2B), beginning with an Aligned session in which pen-tip motion was matched to the physical pen, with additional Pen-tip Localization blocks interleaved with Hand Localization blocks. Each repetition ended off with a Hand No Cursor and No Pen block. Target and arc positions were pseudo-randomized across trials.

Following a short break, participants completed the rotated session, in which the tip of the pen was rotated 30° counterclockwise. Participants were informed that the pen would behave differently but were instructed to continue aiming directly to the target with fast, straight, and accurate movements. As in the Hand Experiment, the first block of Pen-to-Target trials was extended to 100 trials to allow full adaptation. This was followed by the same sequence of Hand/Pen Localization, No Cursor/Pen, and Pen-to-Target trials. Participants wore the headset throughout the experiment, with separate scenes for hand and pen conditions.

#### Procedure

The overall trial structure in the Pen Experiment closely followed that of the Hand Experiment, with the addition of two new trial types: Pen-tip Localization and No Pen reaches. These were interleaved alongside the standard Hand Localization and No Cursor blocks, as illustrated in Figure 5. To prevent confusion when switching between hand and pen trials, participants received clear verbal and on-screen instructions at each transition.

In the Aligned session, participants completed three repetitions of reach and localization blocks that included both hand and pen conditions to establish baseline performance. Each cycle involved Pen-to-Target trials, Hand Localization, Pen-tip Localization, No Cursor (hand), and No Pen (pen-tip without visual feedback) blocks. Participants were given short breaks throughout but kept the VR headset on. After completing the Aligned session (approximately 50 minutes), they received a mandatory break of at least 20 minutes without the headset, with the option to return the next day to reduce fatigue from the extended protocol.

In the Rotated session, the movement of the pen tip was rotated 30° counterclockwise relative to hand movement. To acquire the target, participants had to compensate by reaching 30° clockwise, while also accounting for the spatial offset introduced by holding the pen. The Rotated session followed the same block structure as the Aligned session, but the initial Pen-to-Target block was extended to ensure full adaptation to the perturbation. As in the Hand Experiment, participants were instructed both on-screen and verbally to not use any conscious strategy during the no cursor and no pen trials so that resulting reach aftereffects would reflect implicit adaptation.

After completing the six rotated blocks with the pen, participants switched back to their hand for a final rotated block consisting of Hand-to-Target, Hand Localization, and Hand No Cursor trials. This final sequence was included to assess whether adaptation transferred back to the hand following tool use. Once complete, participants removed the headset, marking the end of the approximately 50-minute session.

### Data analysis

All position data were recorded using Unity 3D (2020.1.17, Unity Technologies, San Francisco, CA). Data preprocessing and statistical analyses were conducted in R version 3.4.4. The full dataset and analysis scripts can be found in this project’s Open Science Framework repository (https://osf.io/vzds5/).

We aimed to assess whether shifts in perceived location of an end-effector, following visuomotor rotation adaptation is also observed in VR environments and whether they extend to hand-held tools. We compared performance across three main experimental groups: hand-cursor 30°, hand-cursor 60° (from the Hand Experiment), and pen-tip 30° (from the Pen Experiment). Within the pen-tip 30° group, two additional trial types were included: hand-during-pen (hand localization trials interleaved during pen training) and hand-after-pen (a final block of hand trials completed after pen training). These allowed us to examine changes in hand representation across tool use. Adaptation performance: Adaptation was assessed using the first 100 Hand/Pen- to-Target trials in the rotated session. Performance was quantified by calculating the angular deviation between the direction of the cursor (or pen tip) at 3 cm from the home position and the direction of the straight line from home to target. This measure captured early, ballistic movement direction, prior to any corrective adjustments.

To measure adaptation, outward reaches during the first 100 training trials of Hand/Pen- to-Target trials in the Rotated session were analyzed. Adaptation performance was quantified as the angular deviation between the cursor direction from home position and the angle of a straight line from home to target when the cursor was at 3 cm out from home to obtain the ballistic part of the movement before correction.

For no-visual-feedback trials and localization trials, we analyzed five data sets: hand-cursor 30° and hand-cursor 60° from the Hand Experiment, and pen-tip 30°, hand-during-pen, and hand-after-pen from the Pen Experiment. For these measures, we calculated the angle of each point relative to the home position (e.g., the endpoint of a reach without visual feedback or a point indicated during localization trials). Estimates of unseen hand or pen location were obtained by subtracting the actual endpoint angle from the indicated position angle, both relative to the home position. Because participants could freely move their unseen hand or pen tip, movements were not restricted to specific angles. The difference between responses in the aligned and rotated sessions reflected the shift in perceived location caused by training with rotated feedback. Adaptation and aftereffects are reported with opposite signs, as adaptation is in the direction opposite to the rotation, whereas localization shifts follow the same direction as the rotation. For statistical analyses, we used median values for each participant to reduce the influence of outliers.

### Learning performance

The rate and extent of adaptation were assessed by comparing the first two and the last set of four trials (one reach to each target, presented in random order) from the first 100 Hand/Pen-to-Target trials in the rotated sessions (see first Hand/Pen-to-Target task in Figure 2). A 3 (1st, 2nd, last set) × 4 mixed-effects ANOVA, with Greenhouse–Geisser corrections applied, was conducted on degrees of error, with group (hand-cursor 30°, hand-cursor 60°, pen-tip 30°, hand-after-pen) as the between-subjects factor. Significant main effects and interactions were explored as described in the Results, with particular attention to the effect of rotation magnitude on hand-cursor adaptation. To examine whether adapting to the perturbed pen conferred any benefit when subsequently adapting to the rotation with the cursor, we conducted a 2 × 2 ANOVA on the first two sets of training trials for the hand-cursor 30° and hand-after-pen conditions with Bonferroni corrections applied to any follow-up paired comparisons.

### Aftereffects and localization

For the main aftereffects and localization analyses, trials ±3 SD from the mean in the no-cursor and no-pen conditions were removed. This excluded 0.88% and 0.25% of trials from the hand-cursor 30° and 60° groups (Hand Experiment), and 0.59% from the Pen Experiment. Median angular reach deviations were calculated for each participant by subtracting the endpoint angle (point where hand/pen tip intersects the arc) from the target angle. We tested for error deviations between the aligned and rotated sessions in the No Cursor and No Pen tasks by running paired t-tests between the aligned and rotated sessions for each group with Bonferroni corrections applied. We then took the difference between these rotated and aligned deviations to compute the aftereffects for each group. A one-way ANOVA of these aftereffects were run with group (hand-cursor 30°, hand-cursor 60°, pen-tip 30°, hand- during-pen, hand-after-pen) as between-subject factor to investigate if there were significant differences between conditions in reach aftereffects, with Bonferroni corrections applied to any follow-up paired comparisons.

We then analyzed shifts in end-effector localization found in the hand and pen localization tasks during adaptation. To confirm significance, paired t-tests were performed between the aligned and rotated sessions in each group with Bonferroni corrections applied. We conducted a one-way ANOVA on the shift in localization from aligned to rotated session (subtracted as above for reach aftereffects) with group (hand-cursor 30°, hand-cursor 60°, pen-tip 30°, hand-during-pen, hand-after-pen) as a between-subject factors to see if there were significant differences between conditions in localization shifts after training and applied Bonferroni corrections.

Lastly, we then explored if shifts in pen and hand localization were related to reach aftereffects by computing Pearson’s correlations between participants’ median deviations in the No Cursor and No Pen reaches from both the Hand and Pen experiments.

## Results

### Learning rate during adaptation

#### Learning performance

Before investigating how the addition of a tool during training changes implicit reach aftereffects and localization, we first confirmed that participants did indeed learn the perturbation by the end of the first 100 reach training trials (Figure 4). During initial trials, the errors resembled the size of the perturbation, while by the end of the training, participants were able to compensate for 75-80% of the visuomotor rotation. This was confirmed by a main effect across the three trials set (F_(1.83, 179.60)_ = 92.50, p < .001, η² = 0.27), in a 3 (first, second, and last trial set) x4 (hand-cursor 30°, hand-cursor 60°, pen-tip 30°, hand-during-pen) mixed-effects ANOVA. However, this learning across trials did not vary across the three groups (no 3x4 interaction: F_(3.67, 179.60)_ = 2.37, p = .059, η² = 0.02), suggesting that all groups adapted to their perturbation in a similar manner. Nonetheless, because of the larger errors (more compensation due to larger rotation) in the hand-cursor 60°group (pink curve) compared to the other two groups (Figure 4), there was a main effect of group (F_(2, 98)_ = 10.86, p < .001, η² = 0.12). This main effect of group effect persisted for the two additional 2x3 ANOVAs comparing pen and hand for 30° rotation, and between the 30° and 60° rotation. This suggests that across training, errors were a bit smaller for the pen group (purple curve) compared to the hand-cursor 30° group (blue curve in middle panel) at least for the trial sets tested. Given there were no interactions between trial and group for either the 3x4 nor follow-up ANOVAs, the overall reduction of errors, and the rate and amount of compensation, was similar across groups.

**Figure 4.**
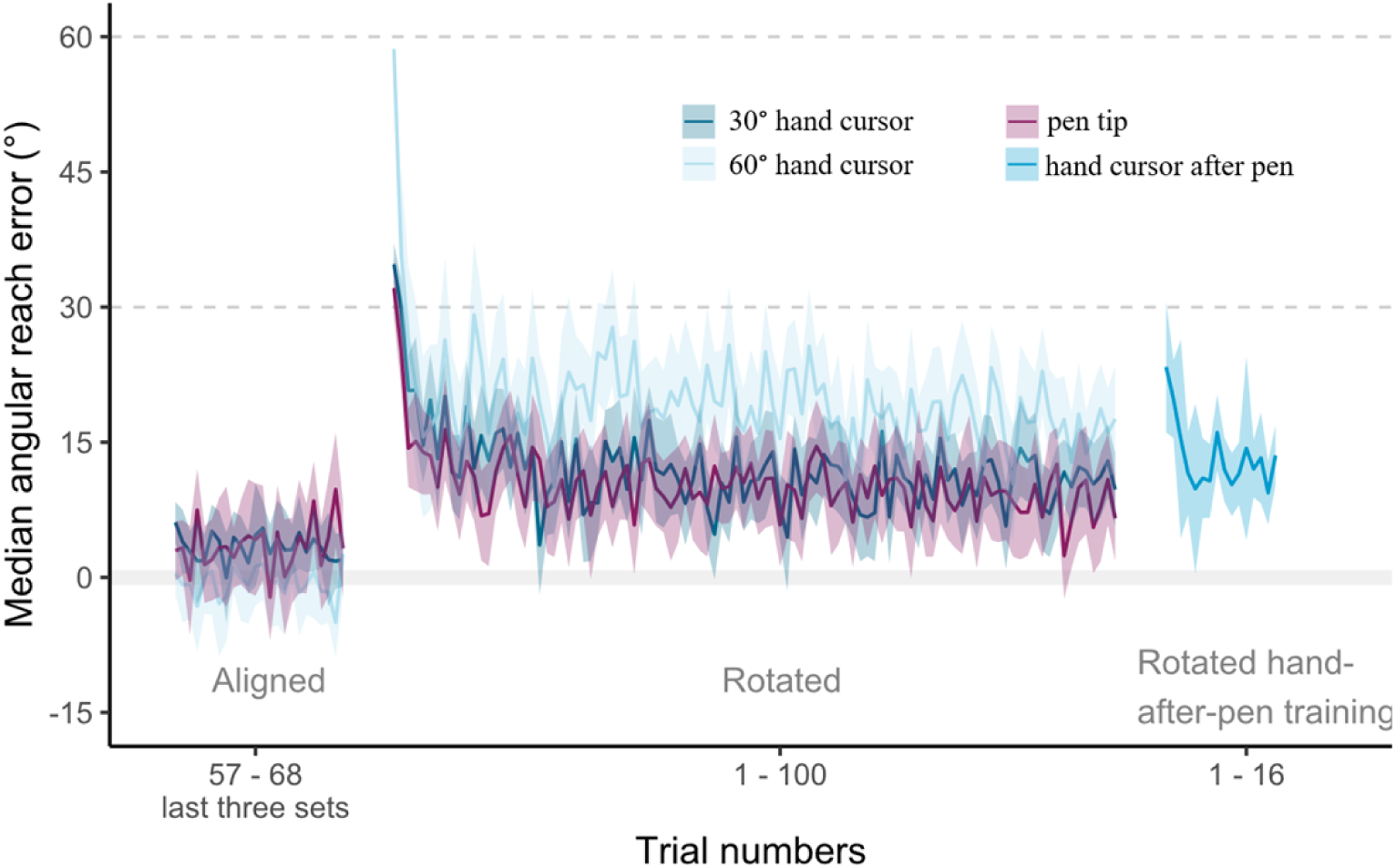
Learning during adaptation training. These plots represent the median angular reach deviation across all participants for each trial when participants reached to the target using their hand for all three groups. The aligned phase depicts the last three sets (24 trials) of the aligned session (baseline data). The rotated phase consists of the first 100 trials of training in the rotated session. Hand after pen consists of the last chunk of Pen Experiment and had 32 trials where participants trained their hand after training with the pen. The grey solid line at the 0° mark indicates full compensate to the perturbation.

To determine whether people adapted faster in the hand-cursor 30° group following adaptation with a pen, we compared adaptation in the hand-after-pen group (rightmost blue curve) to the original hand-cursor 30° group (middle blue curve) for the first two trials (Figure 4). We only tested the first two training trials with the hand, as we expected the generalization effect to be greatest at the start. Consistent with this, errors for the rotated hand-cursor after training with a rotated pen were smaller, indicating greater adaptation by the second trial (t_(122)_ = 2.98, p = .018) but not the first trial (t_(122)_ = 2.21, p = .128). As predicted, adapting to a perturbed pen did facilitate adaptation to the same perturbation applied to the hand cursor.

### Reach aftereffects in hand and tool

We tested whether reach aftereffects emerged in the hand and pen experiments by analyzing reach trials without visual feedback (no-cursor, no-pen) before and after adaptation. Paired t-tests between the aligned and rotated types confirmed that aftereffects were produced for all groups as illustrated in Figure 5A&B (p<0.001 for all five paired comparisons). We compared aftereffect magnitude across groups using a one-way ANOVA (hand-cursor 30°, hand-cursor 60°, pen-tip 30°, hand-during-pen, hand-after-pen; Figure 5C), which showed a significant group effect, F(4,172) = 2.46, p = .047. Follow up tests revealed that the hand-after-pen training group exhibited 4.65° larger aftereffects than the hand-cursor 30° group (*t*_(172)_ = 2.54, p = .048).

**Figure 5.**
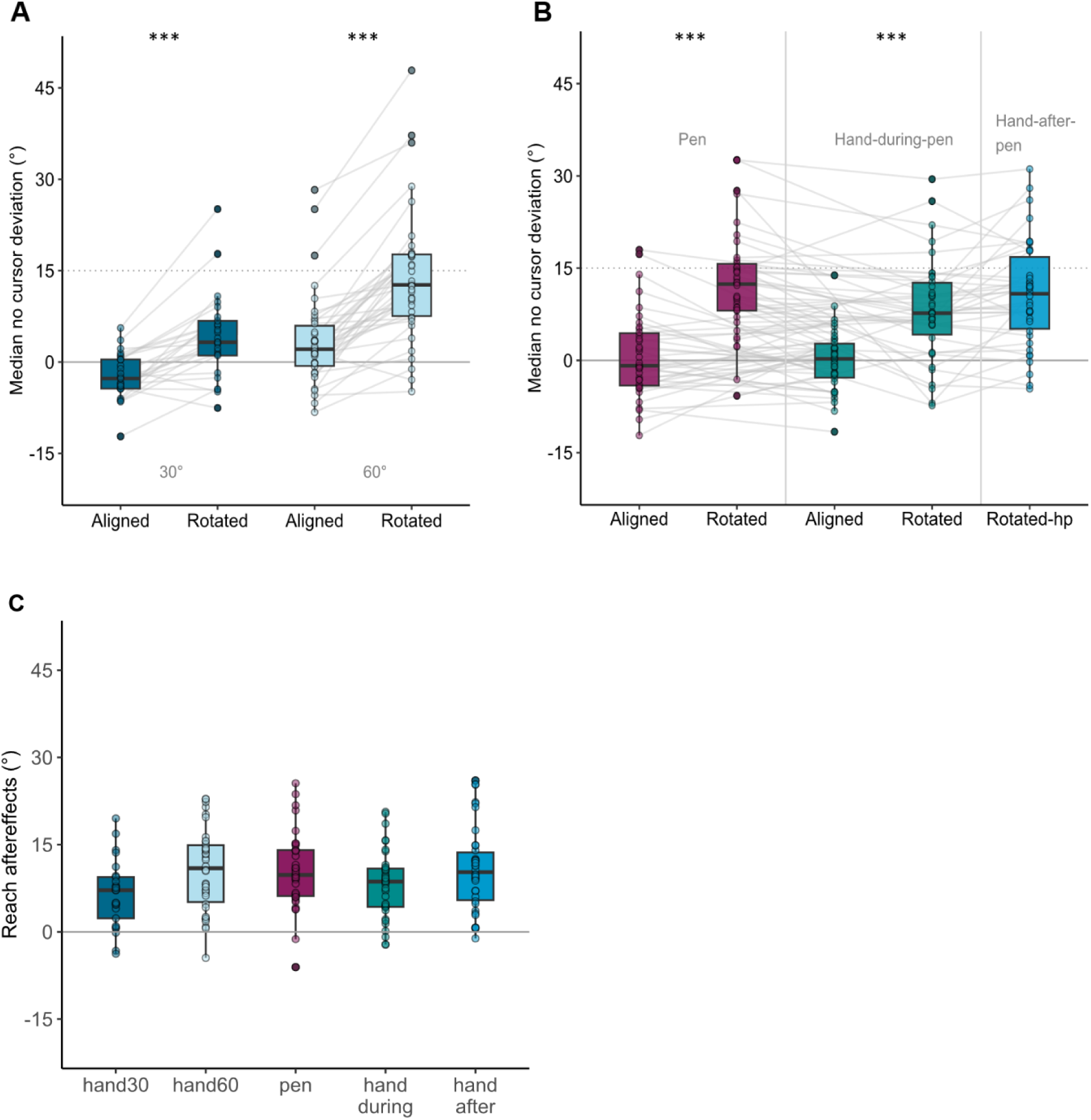
No-cursor/no-pen deviations and aftereffects. The boxes summarize the data distribution, providing information on the median, spread, and presence of outliers across trials. The data was inverted for easier comparison between the no-cursor and localization plots. Each data point represents a single participant’s median reach deviation across all no-cursor/no-pen trials (A) In the Hand Experiment, median reach deviations were plotted against type (aligned and rotated) for each participant for the 30° (left) and 60° (right) groups. (B) shows the median reach deviations in the Pen Experiment for no-pen trials (left), for no-cursor trials during pen training (when participants moved their hand to the target; middle), and no-cursor reaches after subsequently training with the pen (right). (C) illustrates the reach aftereffects across the different groups.

However, no significant differences were found between the hand-cursor 30° and hand-cursor 60° groups (*t*_(172)_ = 2.44, p = .064) nor between the hand-cursor 30° and hand-during-pen (*t*_(172)_ = 1.04, p = 1.00). In addition, reach aftereffects in the hand did not significantly differ during or after the pen training (*t*_(172)_ = 1.68, p = .380). While some group differences reached statistical significance, the magnitude of these effects was small (a few degrees) and may not reflect meaningful differences in practice. Overall, aftereffects emerged after training with both the cursor and with a pen in a virtual environment.

### Shifts in end effector localization

We next examined whether shifts in end-effector localization emerged following training across sessions. Specifically, we assessed localization shifts for the hand in the Hand Experiment (during training with a 30° and 60° rotated hand-cursor) and for the pen and hand in the Pen Experiment (during pen training and hand-after-pen training). As shown in Figure 6, all groups demonstrated significant shifts in localization (p<0.001), except the hand-cursor 30° group.

**Figure 6.**
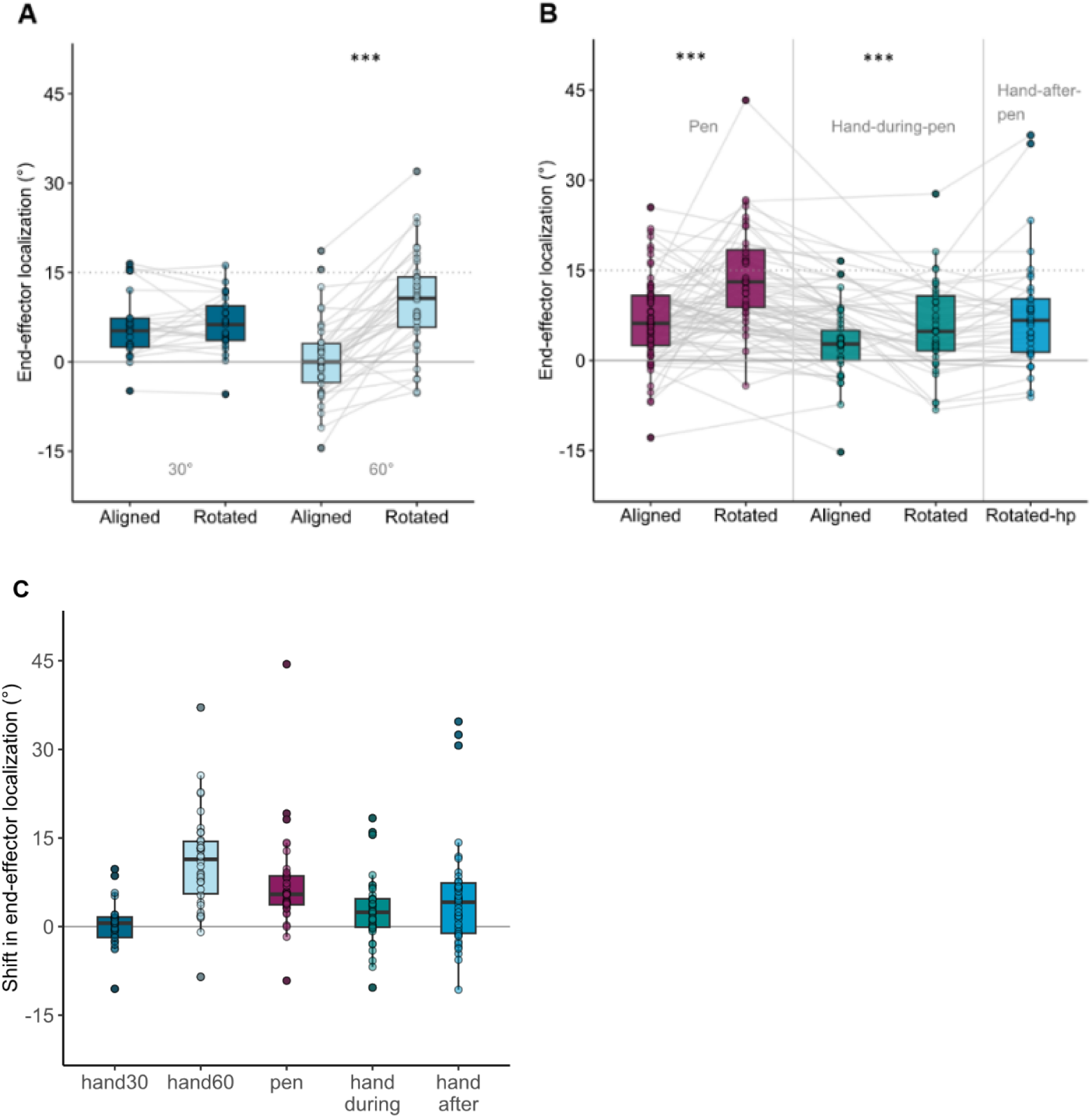
End-effector localization and shifts. The boxes summarize the data distribution, providing information on the median, spread, and presence of outliers across trials. Each data point represents a single participant’s median localization errors relative to the actual position of the hand or pen in degrees angle, across all no-cursor/no-pen trials. (A) In the Hand Experiment, median hand end-effector deviations were plotted against type (aligned and rotated) for each participant for the 30° (left) and 60° (right) groups. (B) shows the median end-effector localization error in the Pen Experiment for pen (left), hand during pen training (middle), and hand after training with the pen (right). (C) illustrates the shifts in end-effector localization in each condition.

A one-way ANOVA on localization shifts (hand-cursor 30°, hand-cursor 60°, pen-tip 30°, hand-during-pen, hand-after-pen) revealed a significant group effect, F(4,172) = 9.71, p < .001. Although the hand-cursor 30° condition did not show a significant aftereffect, the hand-cursor 30° group did not differ from the hand-during-pen (t(172) = 1.18, padj = .965) or hand- after-pen (t(172) = 2.39, padj = .072) groups. But, shifts were larger in hand-cursor 60° than hand-cursor 30° (t(172) = 5.28, padj = .004). Hand-during-pen and hand-after-pen did not differ (t(172) = 1.36, padj = .700) from each other. Overall, VR training with either a cursor or pen produced measurable localization shifts—except in hand-cursor 30°—and pen training alone induced shifts in hand localization, indicating that tool training can generalize to body-based localization.

As expected, localization shifts were generally smaller than reach aftereffects, consistent with prior literature. However, the absence of a significant shift in the hand-cursor 30° group— despite exposure to a same perturbation—was unexpected. Inspection of baseline localization in this group revealed a relatively large positive bias, which may have masked a typically small post-training shift in perceived hand position. Notably, the magnitude of this non-significant shift did not differ significantly from the shifts observed in the hand-during-pen and hand-after-pen groups, suggesting that all three conditions produced similarly small changes in hand localization despite differences in statistical significance.

Lastly, we then explored if shifts in pen and hand localization were related to reach aftereffects (Figure 7). We found that the linear regression model that includes both the main effects of aftereffects and shift as well as their interaction with the group variable was significant (F_(9,167)_ = 10.45, p < .001, r^2^ = 0.326, slope = 0.36). That is, larger reach aftereffects were associated with larger shifts. And while the slopes were all positive for all groups, the relationship for most of the groups was weak with r^2^ of 0.08 to 0.16, except for the hand-cursor 60° group (β = 0.44, r^2^= 0.463, p = .085). Nevertheless, if both reach aftereffects and localization shifts rely on implicit adaptation processes, it makes sense that they are somewhat related

**Figure 7.**
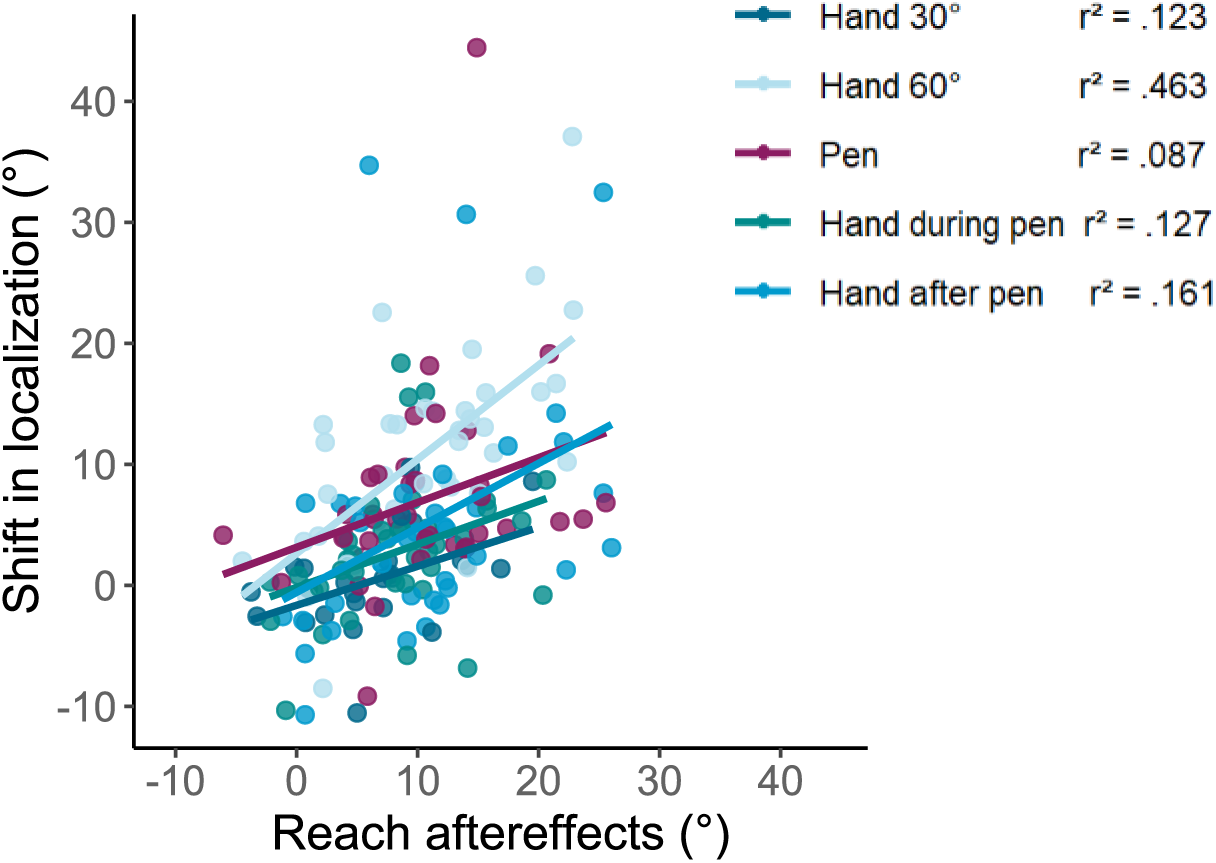
Relationships between changes in hand and pen localization, reach aftereffects for no cursor and no pen reaches across both the Pen and Hand Experiments. The y-axis shows the mean shift in localization for each participant and the x-axis represents each reach aftereffects for both hand and pen for each participant. The solid lines correspond to a regression line and individual data points are color coded according to the corresponding groups.

## Discussion

This study investigated whether adapting to reaching movements using a hand-held tool (a pen) in an immersive virtual environment affects the perceived location of the unseen tool and hand. We also compared these effects to those observed using a conventional hand-cursor interface in VR environment. In the Hand Experiment, participants reached to visual targets with a rotated hand-cursor, either 30° or 60° offset from actual hand movement. While learning rates were comparable across groups, those who trained with the larger 60° perturbation exhibited greater aftereffects and showed the typical shift in perceived location of the unseen hand. In the Pen Experiment, individuals adapted to a 30° rotation using the tip of a virtual pen. The rate and extent of adaptation, along with the resulting reach aftereffects, were comparable—or in some cases slightly greater—than those observed in the hand-cursor conditions. Crucially, adaptation with the pen produced comparable aftereffects and perceived shifts in the location of both the unseen pen and the unseen hand within the same experiment. Oddly, although no shifts in perceived hand location were found following the 30° rotation in the Hand Experiment, such shifts did emerge both during pen adaptation and after brief training with the hand-cursor following pen adaptation. This study investigated whether adapting to reaching movements using a hand-held tool (a pen) in an immersive virtual environment affects the perceived location of the unseen tool and hand. We also compared these effects to those observed using a conventional hand-cursor interface in VR environment. In the Hand Experiment, participants reached to visual targets with a rotated hand-cursor, either 30° or 60° offset from actual hand movement. While learning rates were comparable across groups, those who trained with the larger 60° perturbation exhibited greater aftereffects and showed the typical shift in perceived location of the unseen hand. In the Pen Experiment, individuals adapted to a 30° rotation using the tip of a virtual pen. The rate and extent of adaptation, along with the resulting reach aftereffects, were comparable— or in some cases slightly greater—than those observed in the hand-cursor conditions. Crucially, adaptation with the pen produced comparable aftereffects and perceived shifts in the location of both the unseen pen and the unseen hand within the same experiment. Oddly, although no shifts in perceived hand location were found following the 30° rotation in the Hand Experiment, such shifts did emerge both during pen adaptation and after brief training with the hand-cursor following pen adaptation. These findings suggest that the perceptual changes in end-effector localization—typically observed during hand-movement adaptation to visual perturbations in simple 2D displays —also extend to hand-held tools and more realistic looking environments.

These findings demonstrate that the perceptual changes in end-effector localization— typically observed during hand-movement adaptation to visual perturbations in simple 2D displays—also extend to hand-held tools and immersive, realistic environments. In doing so, we move beyond conventional hand-cursor paradigms to show that reach adaptation and perceptual recalibration occur with a familiar tool in immersive VR. By the end of pen training, participants compensated for the visuomotor rotation by about 75–80%, a level of adaptation comparable to that observed in classic [2,4,9,47–53] and immersive VR [28–31,33] visuomotor rotation tasks.

Notably, participants who first trained with the pen demonstrated slightly faster adaptation when subsequently reaching with the hand-cursor in the "hand-after-pen" session. Although the initial reach error (first set of four trials) was comparable to that of naïve participants (i.e., those who had not undergone pen training), the error on the second set was significantly reduced, indicating more rapid early compensation. Interestingly, despite identical targets and rotation and the use the same controller, simply switching from the pen to the hand-cursor led to an increase in initial error—relative to where participants had left off at the end of pen training (see right side of Figure 4). This incomplete transfer of learning, despite using the same controller to move either the cursor or pen, suggests that the motor system treats the pen and hand-cursor as distinct end-effectors, even when all other task parameters are held constant. Thus, adaptation with the pen does not fully generalize to hand-cursor control. The shift in virtual and physical interface—from holding a pen to using the hand directly—appears to be sufficiently salient to disrupt performance, highlighting the specificity of motor learning and the neural representations underlying different effectors.

One possible explanation for the faster adaptation in the hand-after-pen condition is heightened sensitivity to errors, or an explicit or implicit awareness of the visuomotor transformation required across effectors. However, the comparable magnitude of reach aftereffects following both pen and hand training suggests that most of this adaptation was implicit. Additionally, the observed shift in perceived hand location during pen training may have contributed to the accelerated adaptation in subsequent hand-cursor trials. All hand groups, both before and after pen training, exhibited similar aftereffects, consistent with prior work using hand-cursors alone in 2D environments (e.g. [2,7–9,14,16,47,49,54]) and 3D VR environments [28–31]. This suggests that the brain adapts to changes in sensorimotor feedback even when the hand itself is not directly visible, such as when holding a tool.

Numerous studies have demonstrated that training with a misaligned hand-cursor or within a force field can produce robust changes in the estimated position of the hand [9,11,15,16,20,21,23–25,55–58]. The present findings extend this body of work by showing, for the first time, that similar changes in perceived end-effector location also occur with tool use.

Except for the 30° hand-cursor group, all other hand conditions—both during and after pen training—exhibited localization shifts comparable to those previously observed with direct hand-cursor control.

The absence of a significant shift in the 30° hand-cursor group was unexpected. This result may be explained by a large positive baseline bias, which likely masked a typically small post-training shift (see Figure 7). Although this shift was not statistically significant, it was large enough that it was not reliably different from the significant shift produced when estimating the hand during and after pen-training. Overall, the reach aftereffects and localization shifts observed here support the view that visuomotor adaptation involves updating internal models not only for the hand but also for common hand-held tools.

These findings raise the question of whether the observed updates in the perceived end-effector location reflect true tool embodiment. Previous work has shown that extensive tool use can lead to the tool being integrated into the body schema, altering behavioral, perceptual, and physiological responses to the environment ([35,37,43]). Although, Schone et al. ([39]) reported that expert tool users maintain distinct neural representations for the tool and the hand, with greater differentiation across the visuomotor network. Such representational separation is thought to facilitate rapid switching between hand and tool use by reducing interference and preserving task-specific performance.

In the current study, participants exhibited measurable shifts in the perceived location of both the unseen pen and hand, suggesting that these effectors were represented as distinct entities, while still undergoing concurrent updates to their internal models. This pattern points to a flexible representation in the motor network – capable of accommodating tool use without requiring full embodiment. Such flexibility may allow the motor system to integrate tools into ongoing actions when beneficial yet maintain separate mappings to support rapid switching between effectors. Future work should aim to disentangle the neural mechanisms that govern this balance between tool embodiment and tool-specific categorization. Likewise, the use of styluses or controllers in many visuomotor studies may obscure distinctions between hand- and tool-based adaptation, a limitation that could be mitigated by tracking technologies that do not require hand-held devices.

Lastly, in contrast to previous findings, we observed that the magnitude of the cursor rotation influenced both reach aftereffects and the shifts in perceived hand location. While prior studies have reported similar aftereffects and localization shifts across different perturbation sizes ([9,23,48,49]). However, in our study, participants in the 60° hand-cursor group exhibited larger reach aftereffects and localization shifts than those in the 30° hand-cursor group.

Interestingly, the magnitude of these changes in the 60° hand-cursor group was similar to those observed in the 30° hand group during and after pen training. One possible explanation is that the immersive VR environment alters participants’ perception of feedback reliability. In such environments, participants may implicitly treat the feedback as less trustworthy or less representative of real-world conditions, thereby attenuating proprioceptive recalibration. This suggests that adaptation processes in VR may be influenced not only by the nature of the perturbation but also by the perceived credibility of sensory feedback.

As VR becomes more prevalent in rehabilitation settings due to its accessibility and safety, understanding how motor adaptation functions within this medium is increasingly important. VR offers advantages, such as precise manipulation of visual feedback—for example, altering the position of the virtual cursor instead of the entire workspace. This allows for targeted interventions aimed at recalibrating sensorimotor representations. Such understanding is particularly valuable for individuals with neurological disorders that impair sensorimotor function. Although the cerebellum plays a critical role in motor adaptation, individuals with mild cerebellar ataxia have been shown to exhibit proprioceptive recalibration when trained with gradually introduced perturbations [59].

Taken together, our results demonstrate that pen-based training in immersive VR elicits robust reach aftereffects and shifts in end-effector localization. Moreover, this form of training significantly alters both the estimated location of the hand and subsequent hand-only reach behavior, highlighting its potential utility for sensorimotor rehabilitation and tool-use learning in virtual environments.

## Conclusion

This study demonstrates that adaptation to pen-based reaching movements in immersive VR induces significant reach aftereffects and alters the perceived locations of both the pen and the hand. Future research should aim to enhance ecological validity by allowing participants to manipulate tools in VR as they would in real life, incorporating more naturalistic training paradigms. These findings have important implications for VR-based rehabilitation, suggesting that understanding tool interactions within virtual environments can inform the development of more targeted and effective therapeutic interventions. By leveraging the flexibility of VR to manipulate environmental and tool-related parameters, clinicians could design personalized, engaging rehabilitation programs tailored to individual patient needs and goals.

